# Dissecting allele-specific fungicide resistance mechanisms by heterologous expression of the demethylase inhibitor target gene *Cyp51* in a phytopathogen model

**DOI:** 10.1101/2025.10.16.682737

**Authors:** Katherine G. Zulak, Steven Chang, Kar-Chun Tan, Chala Turo, Richard P. Oliver, Francisco J. Lopez-Ruiz

**Affiliations:** School of Molecular and Life Sciences, Centre for Crop and Disease Management, Curtin University, Bentley, WA, Australia 6102; Biosciences, University of Nottingham, Biosciences LE12 5RD, UK

**Author notes:** Corresponding author E.

**Keywords:** Fungicide resistance, *Pyrenophora teres*, *Cyp51*, demethylase inhibitor, *Parastagonospora nodorum*, azole

## Abstract

**BACKGROUND:** Fungicide resistance is a major concern both in agriculture and clinical disease control. Whilst several mechanisms of resistance have been elucidated, assigning phenotype to genotype is often difficult and reliant on correlations.

Resistance to demethylase inhibitor (DMI) fungicides was recently reported in the economically important filamentous fungal barley (*Hordeum vulgare*) pathogen *Pyrenophora teres* f. *teres* (*Ptt*) in Australia. The target of DMI fungicides is encoded by the *Cyp51* gene family; single allele of *Cyp51B* and two copies of the *Cyp51A* gene^1^. Five *Cyp51A* alleles (*W1-A1, 9193-A1, KO103-A1, W1-A2* and *9193-A2*) were identified in *Ptt* with *KO103-A1* containing the mutation F489L *(F495L)* which correlates with resistance to various DMIs.^1^

**RESULTS:** We replaced the coding region of the native *Cyp51B* gene of the filamentous fungal Dothideomycete wheat pathogen *Parastagonospora nodorum* with each of the five *Ptt Cyp51A* alleles to compare the phenotypic effects of each allele in isolation. The native *Cyp51B* of *P. nodorum* could be functionally replaced by *Cyp51-A1* but not *Cyp51-A2*. Transformants carrying *KO103-A1* exhibited significantly higher gene expression than *9193-A1* and *W1-A1*, suggesting the mechanism of gene regulation lies within the coding sequence and is conserved between *Ptt* and *P. nodorum*. The EC_50_ values of the *KO103-A1* transformants were significantly higher than any other transformants or wild type isolates for metconazole, prochloraz and tebuconazole but lower for epoxiconazole.

**CONCLUSION:** This system permits the functional characterisation of fungicide target genes in an isogenic background that mimics the physiological environment of plant pathogens. We suggest the system will prove useful in dissecting the impact of genetic mutations on a spectrum of fungicides and permit the design of fungal strains for screening active ingredients that may control strains resistant to existing fungicides.

## Introduction

*Pyrenophora teres* f. *teres* (*Ptt*; *P. teres* Drechsler; anamorph *Drechslera teres* [Sacc.] Shoem.) is a narrow-host range necrotrophic fungal pathogen responsible for the barley (*Hordeum vulgare*) disease net form net blotch (NFNB). Net blotch diseases are rapidly increasing in prevalence across nearly all barley growing regions.^2^ Together with spot form net blotch (SFNB; *P. teres* f. *maculata)*, these diseases can cause yield losses ranging from 10-40%, and in severe cases, up to 100%.^2^ No-till farming practices have contributed to the spread of disease in recent decades. In the absence of adequately resistant barley cultivars, fungicides have become the main method of disease control.^3^ Quinone outside inhibitor (QoI), succinate dehydrogenase inhibitor (SDHI) and demethylase inhibitor (DMI) fungicides are widely used on barley crops as seed dressings and foliar treatments to control various diseases.^4^ DMI fungicides are site specific fungicides that interact with the heme iron of the cytochrome P450 sterol 14 α-demethylase (Cyp51), thus disrupting ergosterol biosynthesis and inhibiting growth.^5^ These compounds are widely used in both agriculture and medicine to treat fungal pathogens.

Although fungicides remain an important and effective means of control, resistance has emerged as a major global challenge in agriculture. In filamentous fungi, five main mechanisms of resistance to fungicides have been reported; target site modification where mutations in the target gene(s) reduce the binding affinity of the fungicide to the enzyme, overexpression of the target gene(s), increased fungicide efflux by overexpression of membrane-bound transporter proteins, detoxification of the fungicide^6^, and copy number variation in the target gene.^7^ Resistance in *Ptt* to SDHI fungicides is widespread in Europe and Australia and poses a major concern^8,9^. While mutations associated with resistance to QoI fungicides have been detected, field control has not been compromised for all the actives within this fungicide class.^3,10,11^ Resistance to DMI fungicides was first reported in *P. teres* (form unknown) isolates from New Zealand,^12^ though molecular characterization was limited. More recently, Western Australian *Ptt* isolates were found to harbor both target site mutations and overexpression of the DMI target gene *Cyp51*.^1^

Cyp51 is encoded by a small gene family in fungi. A single *Cyp51B* is carried by all ascomycetes, while several species including *Aspergillus* spp., *Fusarium* spp., *Penicillium digitatum, Rhynchosporium commune*, and *Ptt* carry other paralogs.^1,13,14^ *Ptt* carries one copy of *Cyp51B* and two *Cyp51A* genes, *Cyp51-A1* and *Cyp51-A2*.^1^ A survey of *Ptt* isolates from Western Australia revealed five unique *Cyp51A* alleles: *W1-A1, 9193-A1, KO103-A1, W1-A2* and *9193-A2*.^1^ In species with multiple *Cyp51* paralogs, target site-mediated DMI resistance is associated with non-synonymous mutations and/or overexpression of the *Cyp51A* gene.^1,15,16^ Resistance to DMIs in *Ptt* was correlated with the mutation F489L (corresponding to F495L in the archetype, *A. fumigatus*)^17^ in *Cyp51A*, which was only found in isolates with the *KO103-A1* allele.^1^ Resistance factors (RFs) of the resistant isolate KO103 varied by fungicide, with RFs about 10 for tebuconazole, metconazole, triticonazole, difenoconazole and prochloraz, and RFs around 1 for epoxiconazole, prothioconazole, propiconazole and triadimenol.^1^ Additionally, all three *Cyp51* paralogs, *Cyp51-A1, Cyp51-A2* and *Cyp51B* exhibited an enhanced level of tebuconazole-induced gene expression in KO103 compared to the sensitive strain 9193, despite no promoter changes being found.^1^

Linking a fungicide resistance phenotype to specific genotypic alterations is challenging, especially when using naturally occurring isolates^18^. An ideal approach is to express individual resistance alleles in an isogenic background. Yeast expression systems have been instrumental in studying the impact of *Cyp51* mutations on DMI tolerance.^19-21^ Recently, DMI ligand binding studies and Cyp51 reconstitution assays have been used in conjunction with the expression of individual and pairs of relevant mutations in *Candida albicans* to determine the effects of those changes on DMI sensitivity.^20^ Interestingly, amino acid substitutions that resulted in increased tolerance often reduced catalytic activity.^20^ Also, direct ligand binding studies alone failed to identify small differences between mutant Cyp51 proteins, highlighting the need for a combination of DMI ligand binding and half maximal inhibitory concentration (IC_50_) studies.^20^ However, yeast systems have significant drawbacks due to differences in temperature sensitivity compared to plant pathogenic filamentous fungi.^22^ Cyp51 from the wheat pathogen *Zymoseptoria tritici* failed to demethylate Cyp51 substrates eburicol or lanosterol at 37°C but catalysis was restored at 22°C, unlike *C. albicans*.^23^ To address these limitations, a heterologous expression system was recently developed using the Leotiomycete filamentous fungus *Sclerotinia sclerotiorum* to test the SDHI target genes *SdhB* and *SdhC* from various phytopathogens against several actives from this fungicide group.^24^ The *SdhB* and *SdhC* genes from the phytopathogens *Botrytis cinerea, Blumeriella jaapii* and *Clarireedia jacksonii* were introduced into *S. sclerotiorum* and the radial growth on fungicide-amended media validated previously identified resistance mutations. A laboratory mutant *SdhB* allele from *Monilinia fructicola* also conferred resistance to boscalid, suggesting a potential resistance mechanism in *M. fructicola*.^24^

In this study, we have developed a system to express fungal *Cyp51* genes in the model fungal pathogen *Parastagonospora nodorum*, replacing its sole active *Cyp51* gene *Cyp51B* (SNOG_03702). *P. nodorum*, the causal agent of septoria nodorum blotch on wheat (*Triticum aestivum*) is considered an ideal model system for fungicide research as it is a typical filamentous phytopathogen with accessible genomic tools.^22,25^ We replaced the native *Cyp51B* gene of *P. nodorum* with each of the five *Ptt Cyp51A* alleles to assess the phenotypic impact of each allele in isolation and in the same genomic location. We demonstrate that *P. nodorum Cyp51B* can be functionally replaced by the *Ptt Cyp51A1* alleles but not *Cyp51A2*. Transformants carrying the *KO103-A1* allele exhibited differential resistance to various DMIs. Expression of *Cyp51B* in transformants carrying the *KO103-A1* allele was significantly higher upon treatment with sub-lethal tebuconazole doses than either sensitive allele *9193-A1* or *W1-A1*, suggesting a promoter-independent mechanisms of gene regulation. This work introduces a robust new tool in the study of resistance to DMIs and lays the foundation for expanding functional genomic studies to other fungicide classes and also to those fungal species lacking efficient transformation systems.

## Materials and Methods

### Isolates and Culturing

*P. nodorum* strain SN15 (Table 1) was maintained on V8-PDA agar (150 mL Campbell’s V8 juice L^−1^, 3 g CaCO_3_ L^−1^, 10 g Difco PDA L^−1^ and 15 g agar L^−1^) under a 12 h photoperiod at 21°C. When required, spores were produced by incubating plates at 21°C in 12 h cycles of darkness and near-UV light (Phillips TL 40W/05).^26^ *P. teres* f. *teres* (*Ptt*) strains KO103, 9193 and W1 were maintained on V8-potato-dextrose agar plates and incubated at room temperature under white light for 5 – 7 days.^1^ All isolates used are described in Table 1.

**Table 1:**
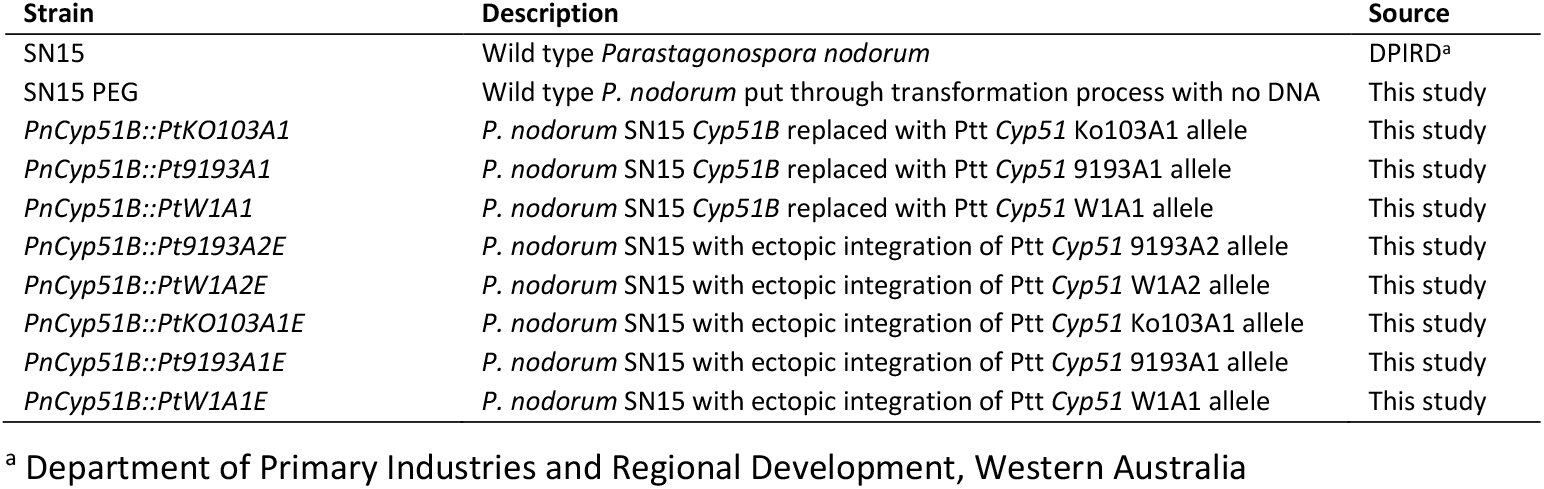
Strains used in this study.

### Construction of the *Ptt Cyp51A* gene replacement vectors and transformation of *P. nodorum*

SN15 strains with the native *Cyp51B* gene (SNOG_03702) replaced by each of the five *Ptt Cyp51A* alleles were generated through genetic transformation with gene replacement vectors constructed using the Gibson assembly® Master Mix (New England Biolabs, Ipswich MA) following the manufacturer’s protocols (Table 1). The gene replacement constructs consist of a 1000 bp flanking region containing the SN15 *Cyp51B* promoter region, *Ptt Cyp51A* cDNA from each of the 5 alleles, a *TrpC* terminator, a *GpdA* promoter, the *hptII* gene that confers resistance to hygromycin B and a downstream 1000 bp flanking region containing the SN15 *Cyp51B* terminator region, all assembled into a pUC18 vector backbone (Figure 1). *Ptt Cyp51* alleles were amplified from *Ptt* isolates 9193, W1 and KO103^1^ using the PttCYP_F and PttCYP_R primers and the remaining construct was amplified from the construct pUC18::TGH3F5F previously made using the TGH3FpUC5F_F and the TGH3FpUC5F_R primers (Table 2). Briefly, this construct was made by amplifying 1000 bp of the native promoter and terminator regions of the *P. nodorum Cyp51B*, and *GpdA, hptII*, and *TrpC* fragments were all amplified from the pAN7-1 plasmid (Table 2, Figure 1). All PCR was performed using Q5 polymerase (New England Biolabs) under the following conditions: 98°C for 30 s, 25 cycles of 98°C 10 s, 64°C for 30 s, and 72°C for 2 min and 30 s followed by extension of 72°C for 2 min. The *P. nodorum* strain SN15 was transformed with each construct using PEG-mediated transformation^26^. Colonies growing through hygromycin overlay were subcultured onto fresh V8-PDA plates containing 200 µg ml^-1^ hygromycin. Presence of the replacement cassette and true gene replacement of the native SN15 *Cyp51B* gene were confirmed using PCR primers SN15_genomic_F and SN15_genomic_R to ensure absence of shorter native SN15 *Cyp51B* gene, and both SN15_genomic_F/CYP51_internal and Hyg-internal_F/Hyg_internal_R to confirm correct integration location (Table 2, Figure 1). True gene replacement events containing a single copy of the cassette were identified using qPCR with the primer sets CN_F1, CN_R1 and CN_F2, CN_F2 (Table 2). The sequence of the integrated *Cyp51A* allele was confirmed by amplification and Sanger sequencing using the PttCYPSeq_F and PttCypSeq_R primers (Table 2, Figure 1). Two isolates containing a single copy integration and one isolate containing an ectopic integration for each *Cyp51A* allele were retained for further analysis. All transformants used in this study are described in Table 1.

**Table 2:**
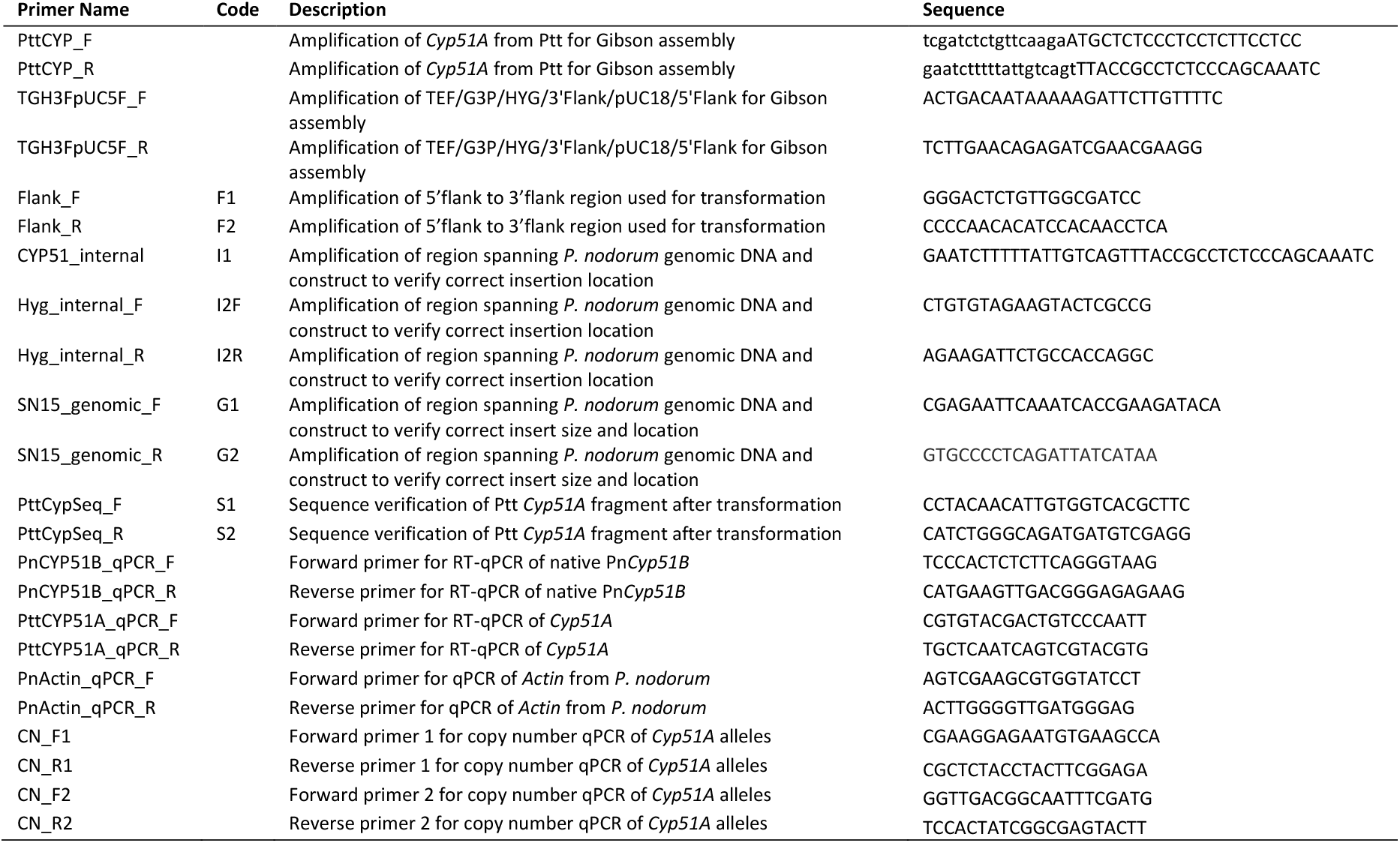
Primers used in this study.

**Figure 1.**
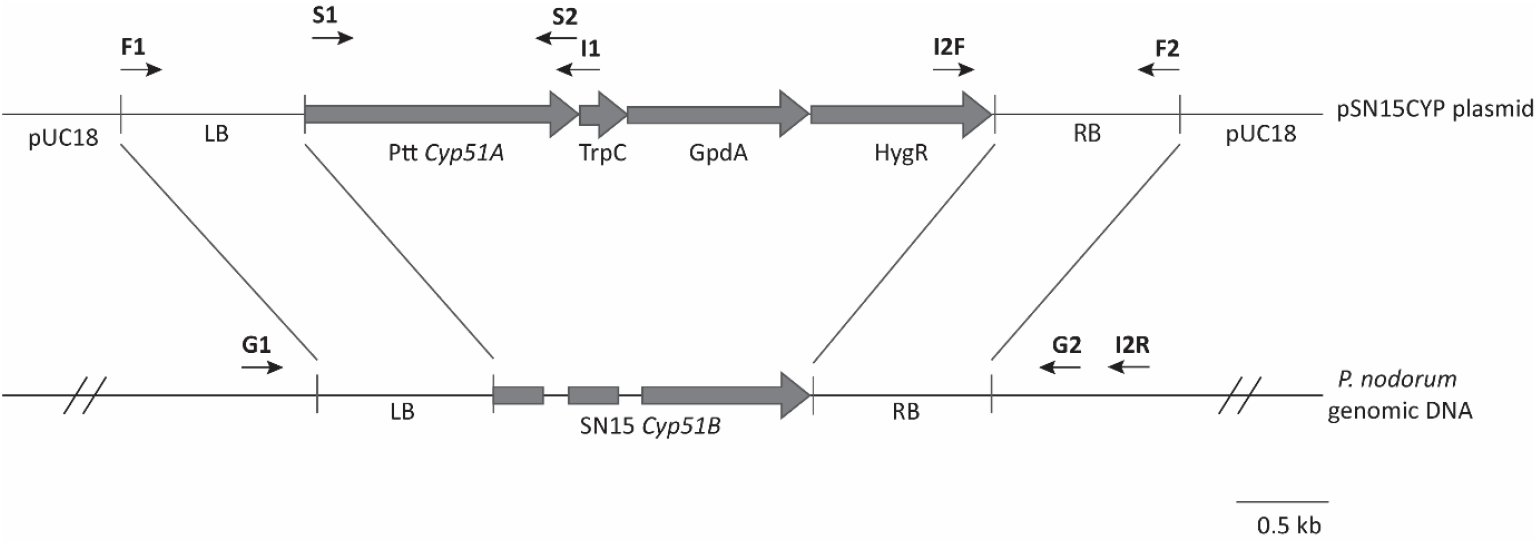
Diagram of construct used for replacement of *Parastagonospora nodorum Cyp51B* with *Cyp51A* allelic variants from *Pyrenophora teres* f. *teres*. Primer codes correspond to primers found in Table 2.

### Gene expression

A final concentration of 1 × 10^6^ spores for each transformant (tebuconazole or ethanol treated control, three replicates per treatment) were grown in 6-well microtitre plates containing 5 ml of Fries2 liquid media,^27^ and incubated at 22°C and 120 rpm. At 72 h post-inoculation, cultures were spiked with either 2 µL of tebuconazole (final concentration of 3.9 µg mL^-1^ for Ko103; 0.31 µg mL^-1^ for W1 and 0.25 µg mL^-1^ for 9193)^1^ or 2 µL ethanol only (the solvent for tebuconazole) and shaken for an additional 24 h. After 24 h, samples were collected, washed twice with sterile DEPC-treated water and extracted with TRIzol Reagent (ThermoFisher Scientific, Waltham MA).^1^ RNA was then treated with Turbo™ DNAase (ThermoFisher Scientific) to remove genomic DNA contamination. cDNA was created using the LunaScript RT SuperMix Kit (New England Biolabs), following the manufacturers’ instructions. qPCR was performed using 2X iTaq Universal SYBR Green Supermix (Bio-Rad Laboratories) and PttCYP51A_qPCR_F/PttCYP51A_qPCR_R and PnCYP51B_qPCR_F/PnCYP51B_qPCR_R primers to measure expression of the inserted *Ptt Cyp51A* gene and the native *P. nodorum Cyp51B* gene, respectively (Table 2). The qPCR reactions were run on a BioRad CFX96 qPCR machine under the following conditions for all genes: 95°C for 5 min, 40 cycles of 95°C for 5 s, and 60°C for 45 s. Melting curve of each primer pairs was analysed after end of the cycle with extension cycle of 65°C to 95°C increment 0.5°C for 5 s. Gene expression of *Ptt Cyp51A* was normalized to SN15 *Act1* (SNOG_01139)^28^ and relative gene expression calculated using the 2^-ΔCT^ method. Each sample was replicated three times.

### Growth rate assays

For each strain, 5 µL of spores (2 × 10^5^ spores mL^-1^) were added to each well of a 96-well microtitre plate containing 95 µL of YSS broth. All transformants and controls were replicated four times and the experiment was repeated twice. Optical density of the plate was measured at 405 nm every six hours for seven days using a Synergy HT microplate reader (ThermoFisher Scientific).

### Fungicide sensitivity assays

A 96-well microtitre plate assay was performed and EC_50_ values calculated for each transformant.^29^ Four technical grade DMI fungicides were used: epoxiconazole, prochloraz, metconazole and tebuconazole (Sigma Aldrich, St. Louis MO). Serial dilutions used for each fungicide were as follows. For epoxiconazole and prochloraz a range of 0, 0.00488, 0.00976, 0.019, 0.039, 0.0781, 0.156, 0.312, 0.625, 1.25, 2.5, and 5 µg mL^-1^ was used, and for metconazole and tebuconazole a range of 0, 0.078, 0.156, 0.313, 0.625, 1.25, 2.5, 5, 10, 20, 40, and 80 µg mL^-1^ was used.

### Detached leaf assays

The first leaf of two-week-old wheat seedlings (var. Wyalkatchem), were cut into 4 cm long pieces and either dipped into water (untreated) or 1.45 µL mL^-1^ Folicur® 430 SC (Bayer CropScience, active ingredient tebuconazole 430 g L^-1^) for 10 s. Treated leaves were then transferred to benzimidazole agar (50 mg benzimidazole, 10 g agar, 1 L Milli-Q water) and dried for about 10 minutes before inoculation. 5 µL spores (10^6^ spores mL^-1^) from each transformant was placed on the leaves to facilitate infection and incubated for 14 days at room temperature under a 12 h photoperiod and then photographed.

### DNA extraction, whole genome sequencing and data analysis

10^6^ spores were incubated in the dark at 22°C, 130 rpm for 7 days. Mycelia were harvested, washed two times with sterile MilliQ water, and freeze dried. High quality genomic DNA was extracted according to Wang, et al.^30^ Library preparation was performed using the Oxford Nanopore Technologies Ligation Sequencing gDNA Native barcoding Kit SQK-LSK109 for the PnCYP51B::Pt9193A1 and PnCYP51B::PtW1A1 transformants and loaded onto a MinION R9 flow cell (Oxford Nanopore Technologies, Oxford, UK). Library preparation was performed using the Oxford Nanopore Technologies Ligation Sequencing gDNA Native barcoding Kit SQK-NBD114.24 and loaded into MinION flow cell R10.4 for the for PnCYP51B::PtKO103A1 transformants (Oxford Nanopore Technologies, Oxford, UK). Resulting data was base-called using Guppy 6.4.6 (high accuracy) for the PnCYP51B::Pt9193A1 and PnCYP51B::PtW1A1 transformants and Dorado 7.6.8 (super accurate) models for PnCYP51B::PtKO103A1 transformants (Oxford Nanopore Technologies, Oxford, UK). The raw reads were corrected and assembled using canu 2.2 with default parameters ^31^. To validate single copy integration, the insert used in the replacement construct (5779 bp) was used as a query to search for the insert in the assembled genome using blastn version 2.9.0 ^32^. A genome with single blastn hit with full length construct confirmed a single integration.

## Results

### Transformation of *Parastagonospora nodorum* with *Ptt Cyp51A* alleles

In order to investigate the role of individual variants of *Ptt Cyp51A* in fungicide resistance, we attempted to replace the native *Cyp51B* gene in *P. nodorum* strain SN15 with each of the five *Cyp51A* alleles found in DMI-resistant and sensitive isolates of *Ptt* (Figure 1).^1^ Single copy correctly replaced transformants were obtained for the *KO103-A1, 9193-A1* and *W1-A1* alleles. In contrast, transformation attempts with *9193-A2* or *W1-A2* alleles yielded only hygromycin-resistant colonies with ectopic integration. Seven attempts were made at the transformation of A2 alleles and over 50 colonies for each attempt were screened for gene replacement. This strongly suggests the A2 variant cannot functionally complement *P. nodorum Cyp51B*.

For transformants containing *KO103-A1, 9193-A1* and *W1-A1*, two single copy, correctly replaced transformants and one single copy ectopic were selected for further analysis. Since single copy correct replacements were not found for *9193-A2* and *W1-A2*, one ectopic transformant was selected for each (Table 1). To corroborate qPCR results and confirm the correct gene replacement, we generated whole genome assembles of two PnCYP51B::PtKO103A1 transformants, and one each of PnCYP51B::Pt9193A1 and PnCYP51B::PtW1A1. A BLASTN analysis using the insert sequence as a query returned a single BLAST hit with over 99.98% identity in all transformants tested. Phenotypic characterisation of transformants showed no significant differences in growth rate among the transformants (Figure 2). Although there were some significant differences in sporulation rate, the results were variable, and no clear pattern could be deduced (Figure 2).

**Figure 2.**
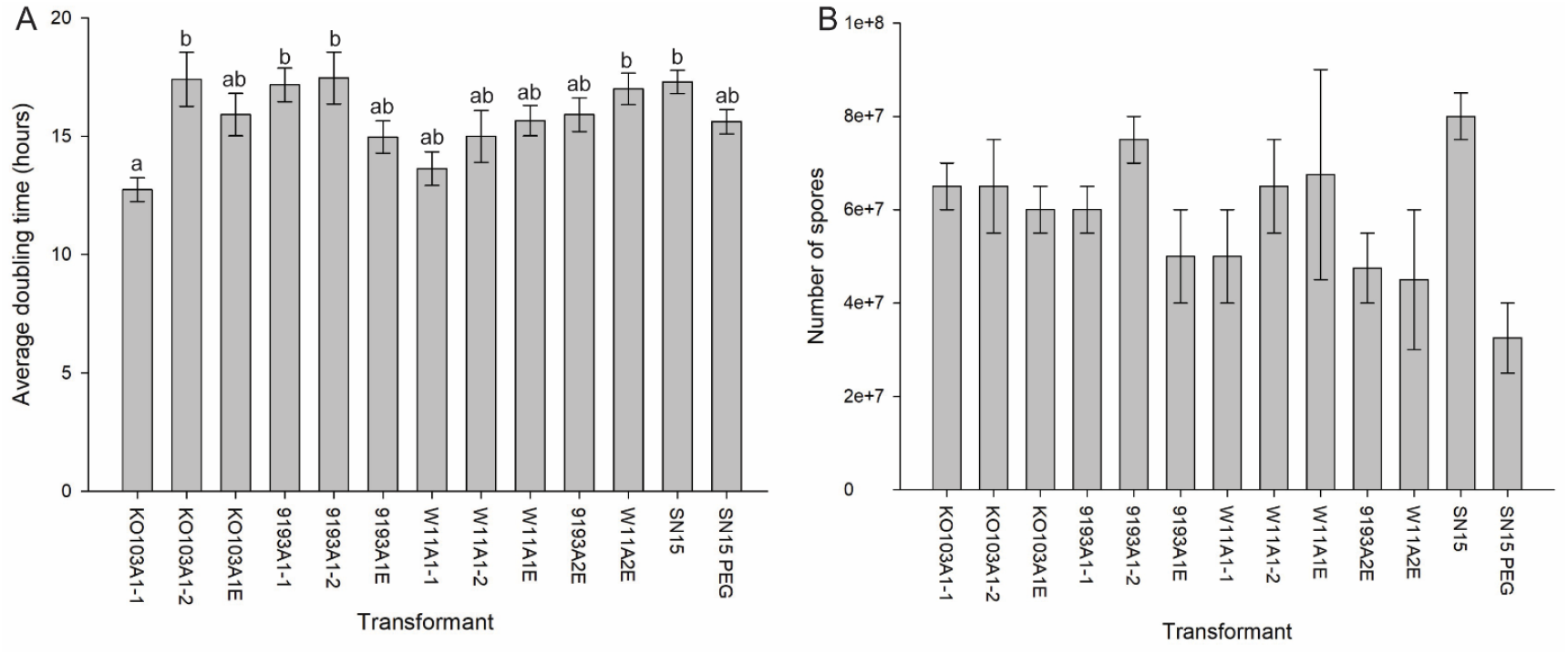
Growth parameters of transformants. A – Average doubling time and B – amount of spores produced for each transformant. N = 4 biological replicates. Strains listed in Table 1.

### Response of transformants to fungicide treatment

To determine the phenotypic impact of each *Cyp51* allele, spore suspensions from each transformant were tested against four DMI fungicides to determine the EC_50_ (Figure 3). Transformants carrying the resistant *KO103-A1* allele (F489L) exhibited significantly higher EC_50_ values than all other transformants or wild-type isolates tested for metconazole, prochloraz, and tebuconazole (Figure 3 B-D). In contrast, when tested against epoxiconazole, EC_50_ values of *PnCYP51B::PtKO103A1* transformants were not statistically different from transformants carrying sensitive alleles *PnCYP51B::Pt9193A1* or *PnCYP51B::PtW1A1* (Figure 3A). Resistance factors for the two *PnCYP51B::PtKO103A1* transformants were also significantly lower (0.85, 0.80) for epoxiconazole compared to metconazole (9.53, 8.46), prochloraz (11.46, 13.19) or tebuconazole (14.01, 12.40; Table 3). These phenotypic responses mirrored those found in the original *Ptt* isolates carrying the different alleles tested in our study.^1^

**Table 3:**
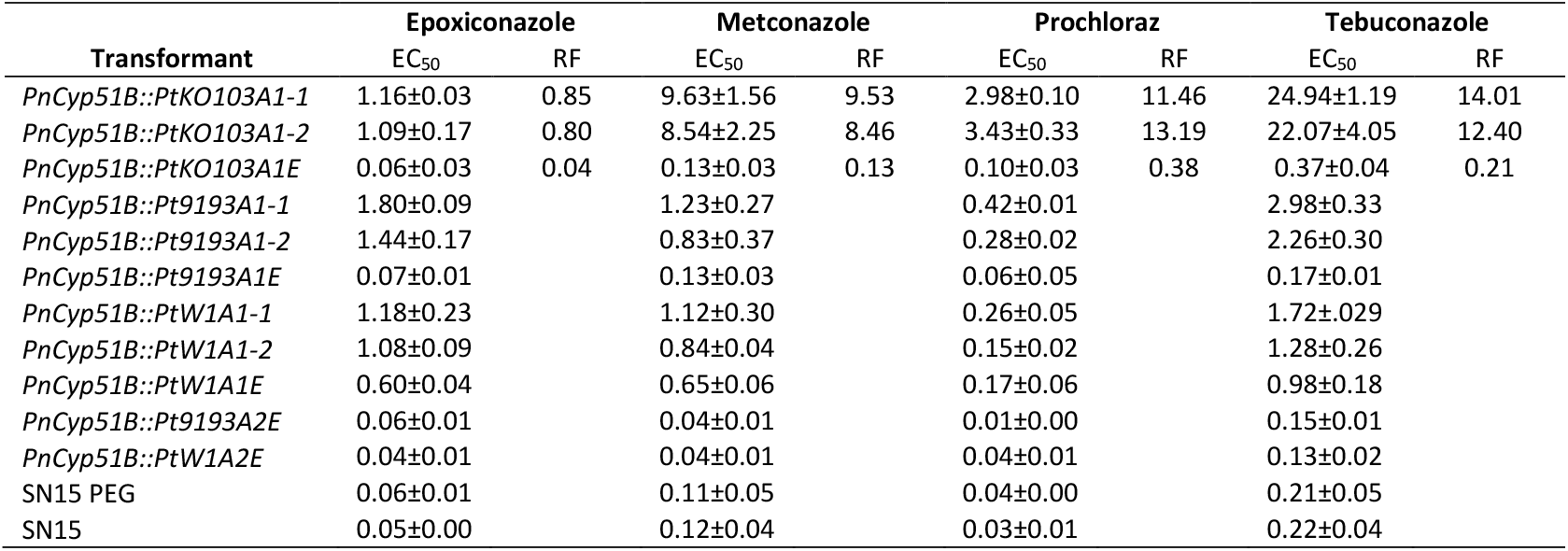
**EC**_**50**_ **and resistance factor values of each transformant treated with epoxiconazole, metconazole, prochloraz and tebuconazole. EC50 values ± standard error of the mean**.

**Figure 3.**
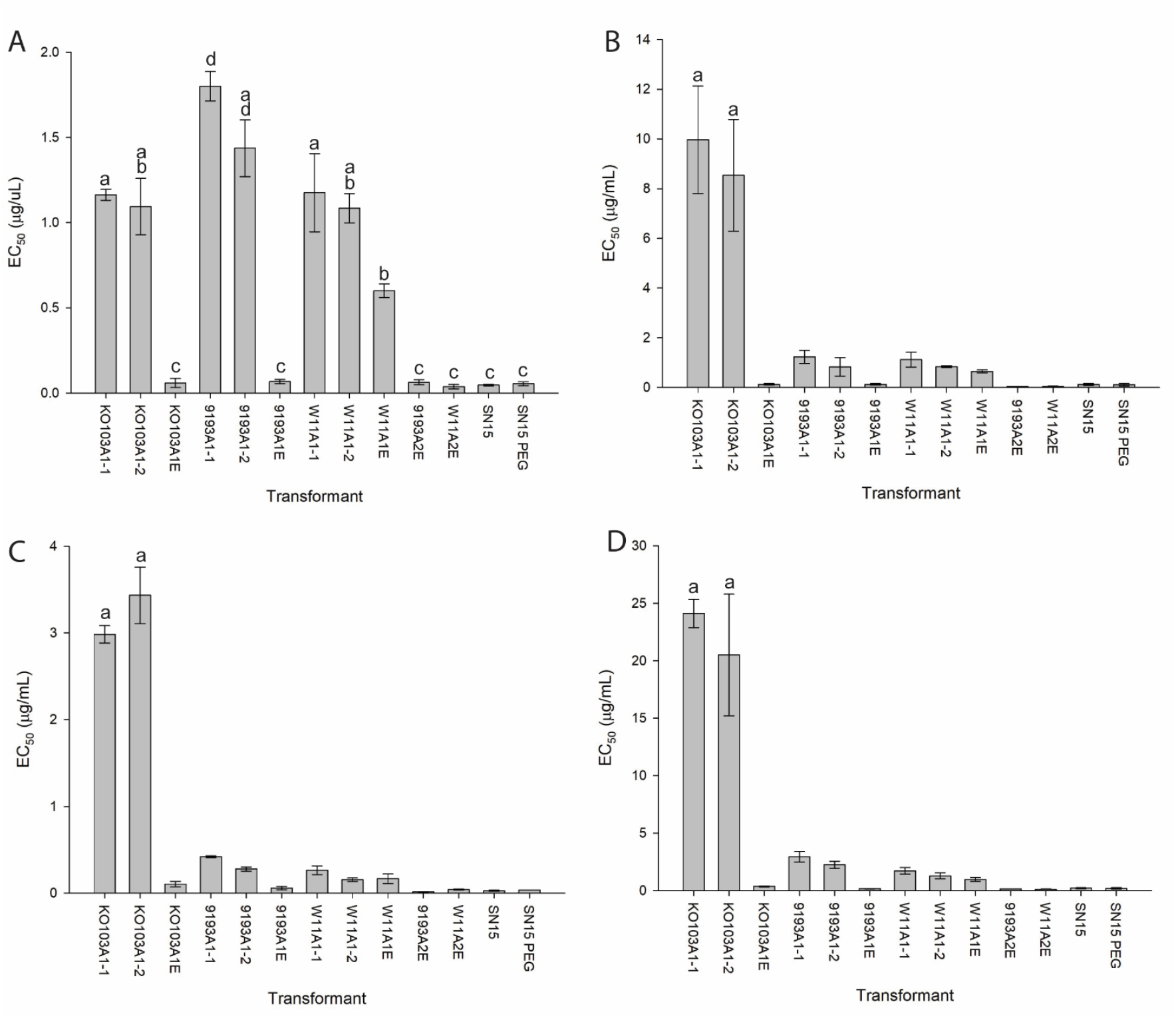
EC_50_ values for all transformants treated with A – epoxiconazole, B – metconazole, C – prochloraz and D – tebuconazole. N = 3 biological replicates per transformant. Letters signify statistical significance p < 0.05 (Tukey). Strains listed in Table 1.

To further validate these findings *in vivo*, each transformant was inoculated onto detached wheat leaves either untreated or dipped in a solution of tebuconazole (Figure 4). Visible lesions formed on untreated leaves except for those inoculated with water. Similar lesions to those found on the untreated leaves were present only in *PnCYP51B::PtKO103A1* transformants and not present in the *PnCYP51B::PtKO103A1* ectopic transformant or in any of the other transformants. Similar to the untreated experiment, the water only control showed no symptoms (Figure 4).

**Figure 4.**
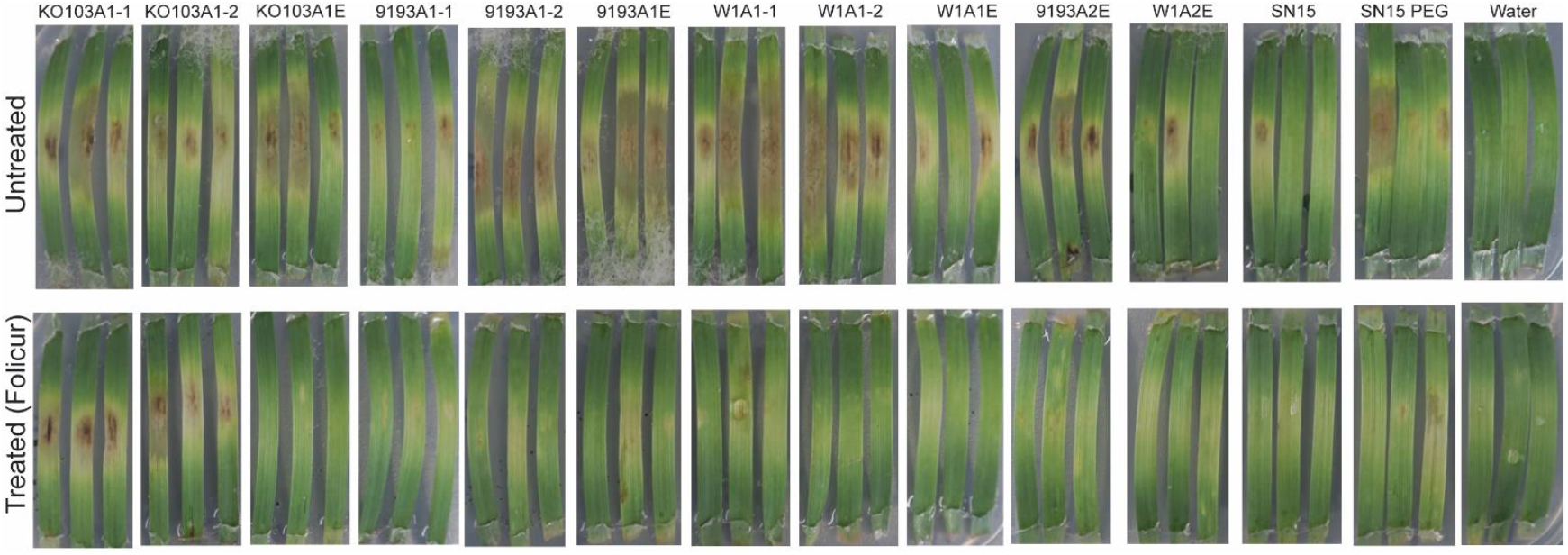
Detached leaf assays of spore suspensions of each transformant on wheat (*Triticum aestivum*, var. Wyalkatchem). Leaves either dipped in Folicur® (430 g L^−1^ tebuconazole) or water (untreated). Photos taken at 14 days past inoculation.

### Gene expression of *Ptt Cyp51A* alleles

Expression levels of *Ptt Cyp51A* were measured in each transformant under untreated and tebuconazole-treated conditions (Figure 5A). A significant induction of *Cyp51A* expression following tebuconazole treatment was observed in the *PnCYP51B::PtKO103A1* transformants. Other transformants exhibited a tebuconazole-related increase, but these changes were not statistically significant. No *Cyp51A* expression was detected in any of the *Cyp51-A1* ectopic transformants, nor in the SN15 PEG control or wildtype SN15. Expression was detected in all the ectopic *Cyp51-A2* transformants but there was no significant difference between control and treated transformants.

**Figure 5.**
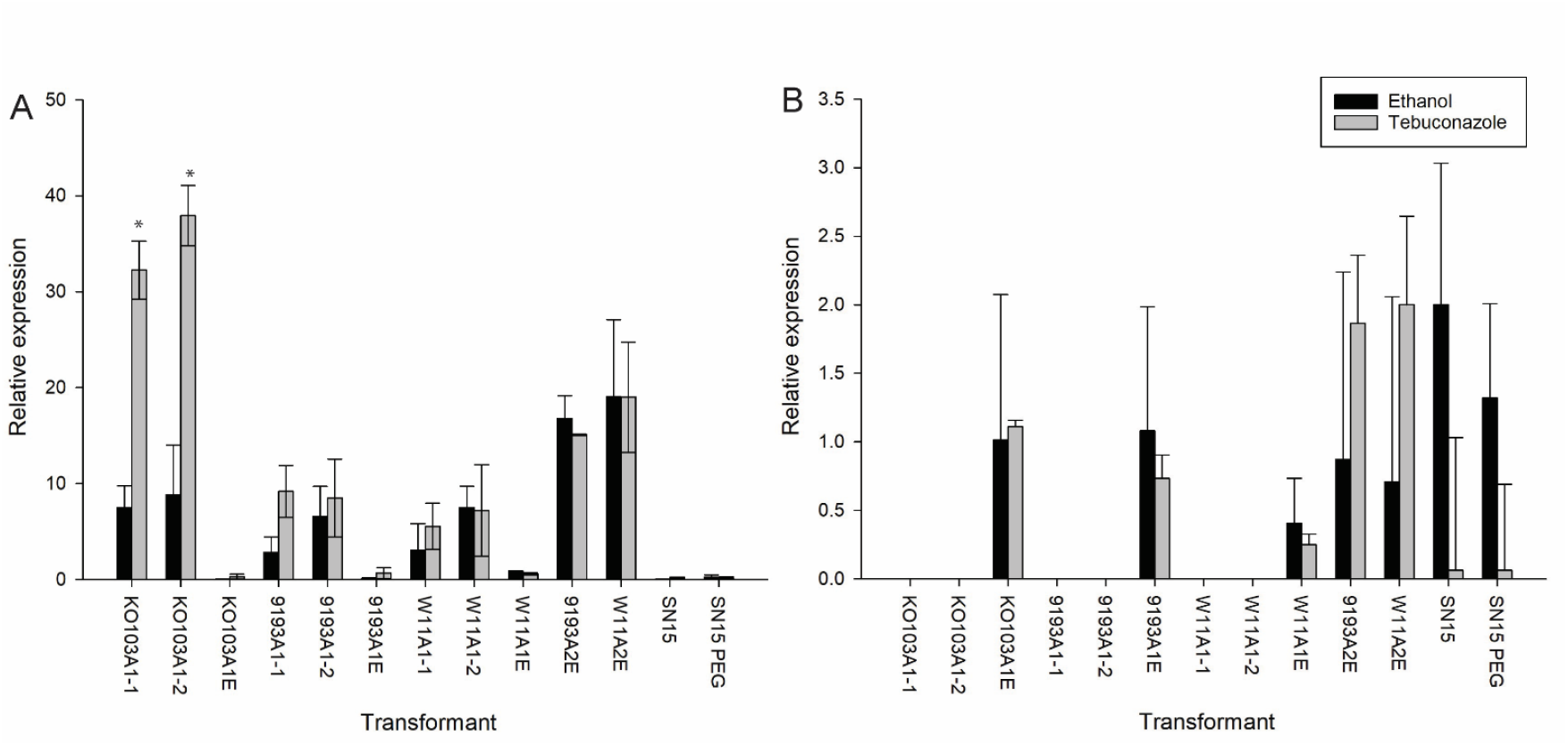
Gene expression analysis of A – *Cyp51A* from *Pyrenophora teres* f. *teres* and B – the endogenous *Cyp51B* from *Parastagonospora nodorum* by qPCR of each transformant either grown in media supplemented with ethanol only (0) or with an EC_50_ relevant to the respective alleles. Mean relative gene expression calculated by 2^-ΔCT^ normalized to Actin. Asterisks signify statistical significance p < 0.05 (Tukey)

Expression levels of the endogenous *P. nodorum Cyp51B* was measured in each transformant as well. No expression was detected in any of the true replacement transformants with the *Cyp51A1* alleles and no statistically significant differences in expression were found for any other transformant (Figure 5B).

## Discussion

In this study, we have developed a heterologous expression system using *P. nodorum* to investigate individual allelic variants of *Cyp51*, the target gene of DMI fungicides. This system allows the isolation of phenotypic effects attributable to individual genetic changes in a gene of interest from changes that may be caused by genomic background variation. As a proof of concept, we expressed the *Cyp51A* gene of *Ptt*, a pathogen currently exhibiting resistance to DMI fungicides in Western Australia.^1^ *P. nodorum* is a well-established model for fungicide research^22,33,34^, offering a complete suite of molecular tools, including reference genome sequences and facile gene replacements techniques.^25^

It is significant that our heterologous system utilises a filamentous fungus rather than a unicellular yeast, providing a more physiologically relevant context for studying fungicide resistance genes in filamentous fungal pathogens. Establishing a direct link between the genotype (allelic variants of *Cyp51A*) and the resulting phenotype (EC_50_ values for various fungicides) is important to dissect the mechanisms of resistance. We successfully replaced the sole functional *Cyp51B* allele *in P. nodorum* with a distantly related *Cyp51A* allele from *Ptt*, despite only 53% amino acid sequence identity (Figure 1, Figure 4). It remains to be seen whether other fungicide target genes such as succinate dehydrogenase (*Sdh*) subunits B, C and D could be similarly replaced in our system.

A recent study using *S. sclerotiorum* as a heterologous transformation system to test impact in SDHI sensitivity of *SdhB* and *SdhC* alleles carrying SDHI-resistance mutations from various phytopathogenic fungi illustrates the utility of such systems.^24^ However, in that study, the endogenous *Sdh* genes were not replaced or silenced, and gene expression of the endogenous and introduced *Sdh* genes was not measured. Thus, it is not known if the introduced alleles exhibited changes in gene expression relative to each other or to the endogenous *Sdh* gene. Consequently, the observed phenotypes likely reflect the added effects of gene sets, rather than introduced allele in isolation.

We found that true replacements of the native *P. nodorum Cyp51B* with *Ptt Cyp51A1* alleles were possible, confirming functional equivalence. This is consistent with a study on *Aspergillus fumigatus* where knocking out either the *Cyp51A* or *Cyp51B* gene did not result in phenotypic differences, suggesting both paralogs could fulfil the same functional role.^35-37^ However, we could not recover a true replacement using either *Ptt Cyp51-A2* alleles. Although gene expression was detected for the A2 alleles (Figure 5), it is possible that the protein products cannot perform the function of the native *P. nodorum* Cyp51B enzyme.

Phenotypic assays revealed that transformants carrying the mutant *KO103-A1* allele exhibited RFs similar to those with wild type *9193-A1* and *W1-A1* alleles when tested against epoxiconazole (average RF = 0.93), but significantly higher for metconazole, prochloraz and tebuconazole (average RF = 12.18, 11.75 and 18.91, respectively; Figure 3, Table 3). These results are consistent with the phenotypes of the original *Ptt* strains carrying the KO103-A1 allele where the RFs for epoxiconazole and tebuconazole were 2.0 and 16.4, respectively.^1^ The EC_50_ results were confirmed *in vivo* for tebuconazole using detached leaf assays, which both corroborates the *in vivo* studies and demonstrates a similar level of pathogenicity of all transformants on leaf tissue (Figure 4).

RNAseq analysis revealed that both *Cyp51-A1* and *Cyp51-A2* alleles in *Ptt* were expressed and upregulated under tebuconazole induction.^1^ In our *P. nodorum* system, all alleles were expressed under the control of the native *P. nodorum Cyp51B* promoter, eliminating promoter variation as a confounding factor. While *Cyp51-A2* alleles were both expressed in ectopic transformants, we cannot confirm translation or protein functionality (Figure 5). There is only one amino acid difference in both *9193-A2* and *W1-A2* that differentiates them from the *Cyp51-A1* alleles (Supplemental Figure S1). Although it is possible that this amino acid change plays an important role in protein stability or enzyme activity, there may be other factors that resulted in the inability to retrieve correctly replaced transformants.

Gene expression analysis of the various transformants showed that only *PnCYP51B::PtKO103A1* transformants exhibited significant upregulation under tebuconazole induction (Figure 5).^1^ Gene expression of tebuconazole treated *PnCYP51B::PtKO103A1* transformants was also significantly higher than any of the other transformants, regardless of treatment (Figure 5). This agrees with the previous finding that *Cyp51A* (no distinction between A1 and A2 alleles) was more highly upregulated in the DMI resistant KO103 isolate than the sensitive 9193 isolate.^1^ This suggests that upregulation of *Cyp51-A1* expression is driven by coding sequence variation rather than promoter differences, and that the regulatory mechanism is conserved between *P. nodorum* and *Ptt*. Also, there was no significant difference in expression of the endogenous *P. nodorum Cyp51B* gene in any of the transformants (Figure 5B). This further suggests that the mechanism observed for the overexpression of the *KO103-A1* allele in *P. nodorum* is promoter independent.

There is only one SNP that differentiates the *KO103-A1* allele from both *9193-A1* and *W1-A1*^*1*^, and that is c1467a, which results in the nonsynonymous mutation F489L^1^ (Supplementary Figure S2). This mutation may affect mRNA stability, as synonymous mutations correlated to changes in gene expression of *Cyp51* have been reported in *Erysiphe necator*.^38,39^ Alternatively, it is possible that the F489L mutation not only interferes with the access of fungicides to the binding cavity of Cyp51, but also affects the binding of its substrate, which may in turn cause a metabolic feedback loop and increased gene expression, perhaps mediated by the accumulation of toxic sterol shunt products.^1^

## Conclusions

This study establishes a heterologous expression system for the analysis of fungal *Cyp51* allele variants in a cytological and physiological settings that is typical for fungal phytopathogens. The system may also be useful to study filamentous animal pathogens. By isolating individual alleles in an isogenic background, we have used the system to gather insight into the function and regulation of *Ptt Cyp51* alleles. Further biochemical and molecular modelling work is needed to fully characterize the impact of specific mutations on both the catalytic function and azole interaction of the allelic variants. The ability to dissect allele-specific effects has proven to be a powerful platform for investigating fungicide resistance and has significant potential for the analysis of other target genes. This study proposes the use of a *P. nodorum* strain for the fine analysis of the impact of individual mutations affecting DMI sensitivity, with broader implications for understanding resistance in both animal and clinical fungal pathogens. Further studies should address the feasibility of replacing other important fungicide target genes, such as the SDH subunits, allowing the simultaneous analysis of multiple resistance mechanisms.

## Acknowledgements

The authors would like to thank Dr. Madeleine Tucker for preliminary work on this system. This study was supported by Curtin University and the Grains Research and Development Corporation through research grants CUR00016 and CUR00023.

## Supplementary Information

### Supplementary Figures

**Supplementary Figure S1.**
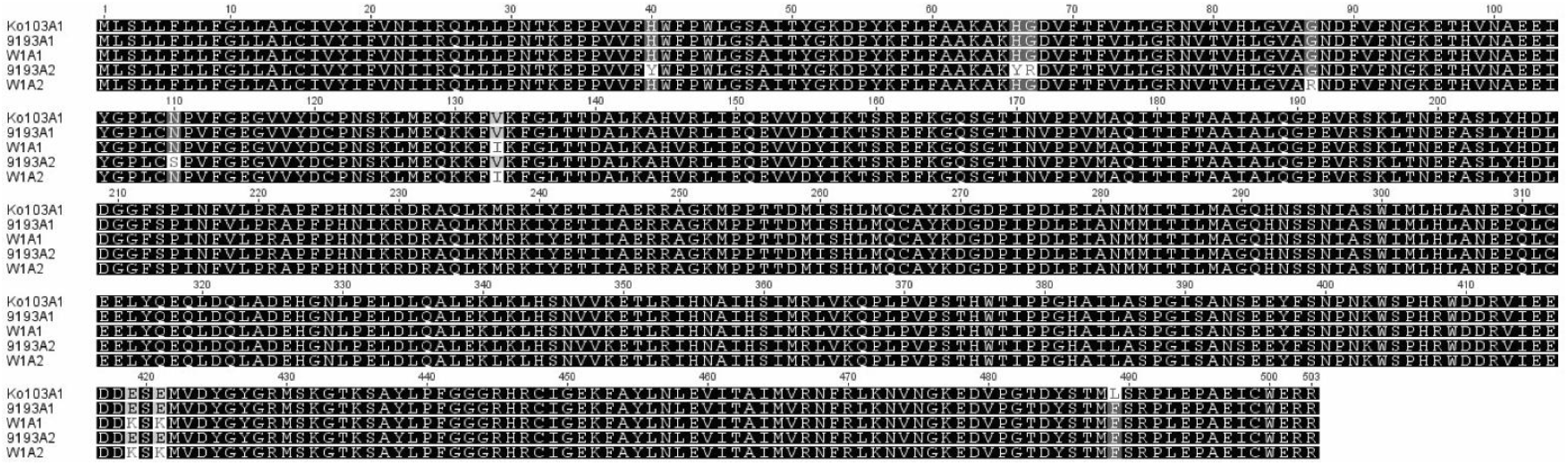
Amino acid alignment of 5 *Pyrenophora teres* f. *teres* allelic variants of *Cyp51A*. Amino acid variations are shaded in grey or white.

**Supplementary Figure S2.**
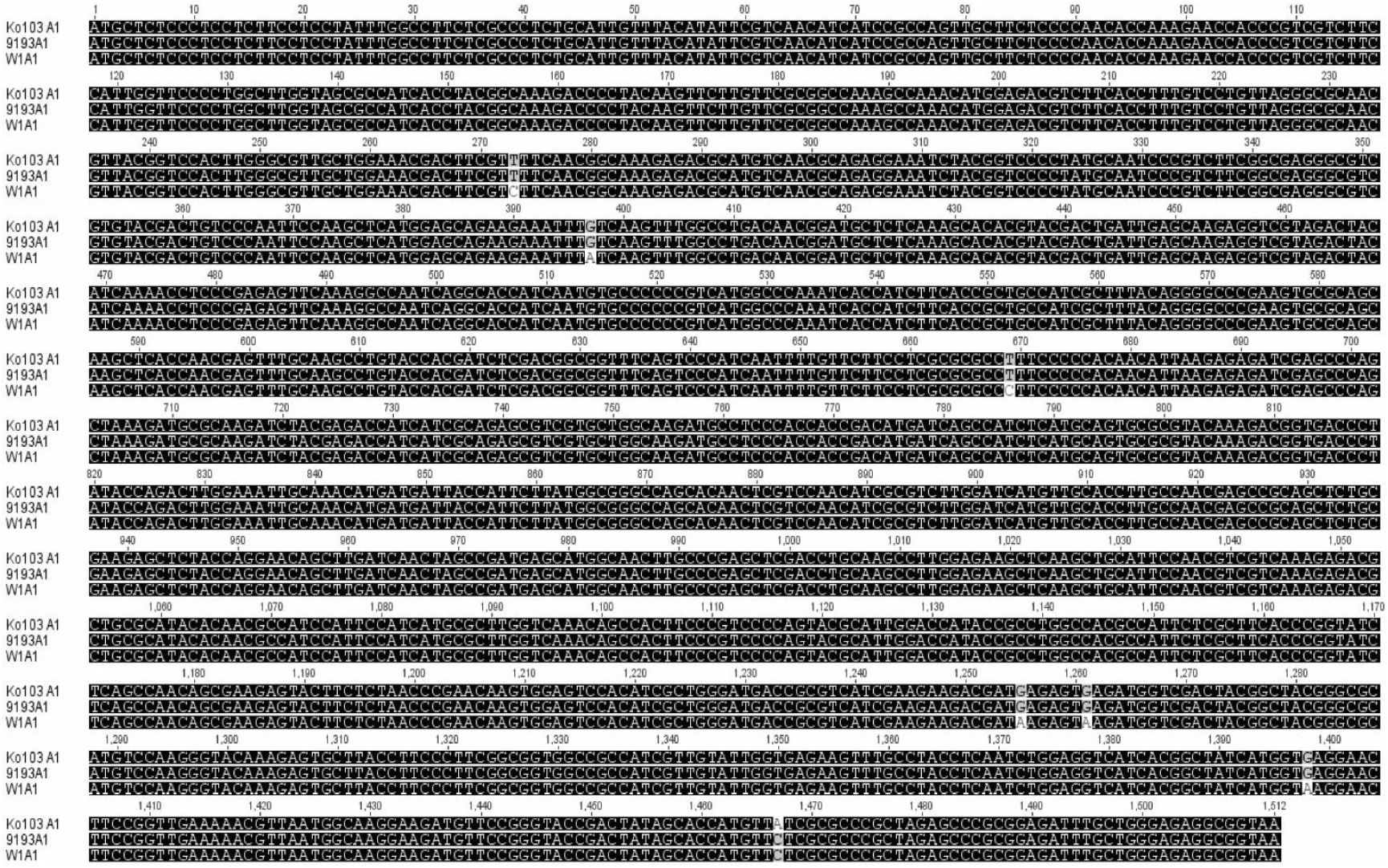
Nucleic acid alignment of *Pyrenophora teres* f. *teres Cyp51-A1* alleles. Nucleic acid differences are shaded in grey or white.

## References

1 Mair, W. J. et al. Demethylase inhibitor fungicide resistance in Pyrenophora teres f. sp. teres associated with target site modification and inducible overexpression of Cyp51. Front. Microbiol. 7, 1279–1296 (2016). 10.3389/fmicb.2016.01279

2 Liu, Z., Ellwood, S. R., Oliver, R. P. & Friesen, T. L. Pyrenophora teres: profile of an increasingly damaging barley pathogen. Mol. Plant Pathol. 12, 1–19 (2011). 10.1111/j.1364-3703.2010.00649.x

3 Rehfus, A. et al. Emergence of succinate dehydrogenase inhibitor resistance of Pyrenophora teres in Europe. Pest Manag. Sci. 72, 1977–1988 (2016). 10.1002/ps.4244

4 (APVMA), A. P. a. V. M. A. PubCRIS: Public chemical registration information system search, <https://portal.apvma.gov.au/pubcris> (2016).

5 Yoshida, Y. & Aoyama, Y. Interaction of azole antifungal agents with cytochrome P-45014DM purified from Saccharomyces cerevisiae microsomes. Biochemical pharmacology 36, 229–235 (1987).

6 Lucas, J. A., Hawkins, N. J. & Fraaije, B. A. The evolution of fungicide resistance. Adv. Appl. Microbiol. 90, 29–92 (2015). 10.1016/bs.aambs.2014.09.001

7 Jones, L. et al. Adaptive genomic structural variation in the grape powdery mildew pathogen, Erysiphe necator. BMC genomics 15, 1081 (2014). 10.1186/1471-2164-15-1081

8 Mair, W. J. et al. Emergence of resistance to succinate dehydrogenase inhibitor fungicides in Pyrenophora teres f. teres and P. teres f. maculata in Australia. bioRxiv, 2023.2004.2023.537974 (2023). 10.1101/2023.04.23.537974

9 Rehfus, A. et al. Emergence of succinate dehydrogenase inhibitor resistance of Pyrenophora teres in Europe. Pest Manag. Sci. 72, 1977–1988 (2016). 10.1002/ps.4244

10 Sierotzki, H. et al. Cytochrome b gene sequence and structure of Pyrenophora teres and P. tritici-repentis and implications for QoI resistance. Pest management science 63, 225–233 (2007). 10.1002/ps.1330

11 Semar, M., Strobel, D., Koch, A., Klappach, K. & Stammler, G. Field efficacy of pyraclostrobin against populations of Pyrenophora teres containing the F129L mutation in the cytochrome b gene. J. Plant. Dis. Prot. 114, 117–119 (2007).

12 Sheridan, J. E., Grbavac, N. & Sheridan, M. H. Triadimenol insensitivity in Pyrenophora teres. Transactions of the British Mycological Society 85, 338–341 (1985). 10.1016/S0007-1536(85)80198-4

13 Becher, R., Weihmann, F., Deising, H. B. & Wirsel, S. G. Development of a novel multiplex DNA microarray for Fusarium graminearum and analysis of azole fungicide responses. BMC Genom. 12, 52–69 (2011). 10.1186/1471-2164-12-52

14 Hawkins, N. J. et al. Paralog re-emergence: a novel, historically contingent mechanism in the evolution of antimicrobial resistance. Molecular biology and evolution 31, 1793–1802 (2014). 10.1093/molbev/msu134

15 Fan, J. et al. Characterization of the sterol 14α-demethylases of Fusarium graminearum identifies a novel genus-specific CYP51 function. New Phytol. 198, 821–835 (2013). 10.1111/nph.12193

16 Brunner, P. C., Stefansson, T. S., Fountaine, J., Richina, V. & McDonald, B. A. A Global Analysis of CYP51 Diversity and Azole Sensitivity in Rhynchosporium commune. Phytopathology 106, 355–361 (2016). 10.1094/phyto-07-15-0158-r

17 Mair, W. et al. Proposal for a unified nomenclature for target-site mutations associated with resistance to fungicides. Pest management science 72, 1449–1459 (2016). 10.1002/ps.4301

18 Oliver, R. P. & Beckerman, J. Fungicides in Practice. (CAB Interational, 2022).

19 Cools, H. J. et al. Heterologous expression of mutated eburicol 14alpha-demethylase (CYP51) proteins of Mycosphaerella graminicola to assess effects on azole fungicide sensitivity and intrinsic protein function. Appl. Environ. Microbiol. 76, 2866–2872 (2010). 10.1128/aem.02158-09

20 Warrilow, A. G. et al. The Evolution of Azole Resistance in <em>Candida albicans</em> Sterol 14α-Demethylase (CYP51) through Incremental Amino Acid Substitutions. Antimicrobial agents and chemotherapy 63, e02586–02518 (2019). 10.1128/aac.02586-18

21 Flowers, S. A., Colón, B., Whaley, S. G., Schuler, M. A. & Rogers, P. D. Contribution of clinically derived mutations in ERG11 to azole resistance in Candida albicans. Antimicrobial agents and chemotherapy 59, 450–460 (2015). 10.1128/aac.03470-14

22 Dancer, J., Daniels, A., Cooley, N. & Forster, S. in Septoria on Cereals: A Study of Pathsystems (eds J. A. Lucas, P. Bowyer, & A.M. Anderson) 316–331 (CABI Publishing, 1999).

23 Price, C. L. et al. Novel substrate specificity and temperature-sensitive activity of Mycosphaerella graminicola CYP51 supported by the native NADPH cytochrome P450 reductase. Appl. Environ. Microbiol. 81, 3379–3386 (2015). 10.1128/aem.03965-14

24 Peng, J. et al. A method for the examination of SDHI fungicide resistance mechanisms in phytopathogenic fungi using a heterologous expression system in Sclerotinia sclerotiorum. Phytopathology 0, 0 (2020). 10.1094/phyto-09-20-0421-r

25 Oliver, R. P., Friesen, T. L., Faris, J. D. & Solomon, P. S. Stagonospora nodorum: from pathology to genomics and host resistance. Annu. Rev. Phytopathol. 50, 23–43 (2012). 10.1146/annurev-phyto-081211-173019

26 Solomon, P. S., Tan, K. C., Sanchez, P., Cooper, R. M. & Oliver, R. P. The disruption of a Galpha subunit sheds new light on the pathogenicity of Stagonospora nodorum on wheat. Mol. Plant-Microbe Interact. 17, 456–466 (2004). 10.1094/mpmi.2004.17.5.456

27 Fries, N. Über die bedeutung von wuchsstoffen für das wachstum verschiedener Pilze. Symboo. Botan. Upsalienses 3, 188 (1938).

28 Tan, K. C. et al. A signaling-regulated, short-chain dehydrogenase of Stagonospora nodorum regulates asexual development. Eukaryotic Cell 7, 1916–1929 (2008). 10.1128/ec.00237-08

29 John, E. et al. Dissecting the role of histidine kinase and HOG1 mitogen-activated protein kinase signalling in stress tolerance and pathogenicity of Parastagonospora nodorum on wheat. Microbiology 162, 1023–1036 (2016). 10.1099/mic.0.000280

30 Wang, C., Milgate, A. W., Solomon, P. S. & McDonald, M. C. The identification of a transposon affecting the asexual reproduction of the wheat pathogen Zymoseptoria tritici. Mol. Plant Pathol. 22, 800–816 (2021). 10.1111/mpp.13064

31 Koren, S. et al. Canu: scalable and accurate long-read assembly via adaptive k-mer weighting and repeat separation. Genome Res. 27, 722–736 (2017). 10.1101/gr.215087.116

32 Altschul, S. F., Gish, W., Miller, W., Myers, E. W. & Lipman, D. J. Basic local alignment search tool. J. Mol. Biol. 215, 403–410 (1990). 10.1016/s0022-2836(05)80360-2

33 Li, W. et al. Malayamycin, a new streptomycete antifungal compound, specifically inhibits sporulation of Stagonospora nodorum (Berk) Castell and Germano, the cause of wheat glume blotch disease. Pest Manag. Sci. 64, 1294–1302 (2008). 10.1002/ps.1632

34 Cooley, R. N., van Gorcom, R. F. M., van den Hondel, C. A. M. J. J. & Caten, C. E. Isolation of a benomyl-resistant allele of the β-tubulin gene from Septoria nodorum and its use as a dominant selectable marker. J. Gen. Microbiol. 137, 2085–2091 (1991).

35 Mellado, E. et al. Targeted gene disruption of the 14-alpha sterol demethylase (cyp51A) in Aspergillus fumigatus and its role in azole drug susceptibility. Antimicrob. Agents Chemother. 49, 2536–2538 (2005). 10.1128/aac.49.6.2536-2538.2005

36 Martel, C. M. et al. Complementation of a Saccharomyces cerevisiae ERG11/CYP51 (sterol 14alpha-demethylase) doxycycline-regulated mutant and screening of the azole sensitivity of Aspergillus fumigatus isoenzymes CYP51A and CYP51B. Antimicrob Agents Chemother 54, 4920–4923 (2010). 10.1128/aac.00349-10

37 Garcia-Effron, G. et al. Differences in interactions between azole drugs related to modifications in the 14-alpha sterol demethylase gene (cyp51A) of Aspergillus fumigatus. Antimicrob. Agents Chemother. 49, 2119–2121 (2005). 10.1128/aac.49.5.2119-2121.2005

38 Rallos, L. E. E. & Baudoin, A. B. Co-occurrence of two allelic variants of CYP51 in Erysiphe necator and their correlation with over-expression for DMI resistance. PLoS One 11, e0148025 (2016). 10.1371/journal.pone.0148025

39 Frenkel, O., Cadle-Davidson, L., Wilcox, W. F. & Milgroom, M. G. Mechanisms of resistance to an azole fungicide in the grapevine powdery mildew fungus, Erysiphe necator. Phytopathology 105, 370–377 (2015). 10.1094/phyto-07-14-0202-r

